# Cryo-EM ligand building using generative AI and molecular dynamics

**DOI:** 10.1101/2025.02.10.637508

**Authors:** Nandan Haloi, Rebecca J. Howard, Erik Lindahl

## Abstract

Resolving protein-ligand interactions in atomic detail is key to understanding how small molecules regulate macromolecular function. Although recent breakthroughs in cryogenic electron microscopy (cryo-EM) have enabled high-quality reconstruction of numerous complex biomolecules, the resolution of bound ligands is often relatively poor. Furthermore, automated methods for building and refining molecular models into cryo-EM maps have largely focused on proteins and may not be optimized for the diverse properties of small-molecule ligands. Here, we present an approach that integrates generative artificial intelligence (AI) with cryo-EM density-guided simulations to fit ligands into experimental maps. Using three inputs: 1) a protein amino acid sequence, 2) a ligand specification, and 3) an experimental cryo-EM map, we validated our approach on a set of biomedically relevant protein-ligand complexes including kinases, GPCRs, and solute transporters, none of which were present in the AI training data. In cases for which generative AI was not sufficient to predict experimental poses outright, integration of flexible fitting into molecular dynamics simulations improved ligand model-to-map cross-correlation relative to the deposited structure from 40–71% to 82–95%. This work offers a straightforward template for integrating generative AI and density-guided simulations to automate model building in cryo-EM maps of ligand-protein complexes, with potential applications for characterization and design of novel modulators and drugs.

## Introduction

Protein-ligand binding is of key importance to both biomolecular regulation and pharmaceutical activity. Resolving protein binding of a ligand at the atomic level, including its geometry, pose, and interactions with specific amino acid residues, can critically aid the characterization of endogenous modulators or design of novel drugs. Recent advances in single-particle cryogenic electron microscopy (cryo-EM) have enabled the determination of diverse macromolecular structures, including targets inaccessible by X-ray crystallography, and often with ligands bound [1]. However, even with high-quality data, the resolution of bound ligands is often lower than the surrounding protein, and may be too poor for definitive model building. For example, in a recently reconstructed cryo-EM map for *β*-galactosidase bound to the inhibitor phenylethyl *β*-D-thiogalactopyranoside, the protein was resolved to 1.5 Å, while the ligand densities were limited to 3–3.5 Å [2].

Several automated methods have been developed to refine molecular models in cryo-EM maps, broadly categorized as rigid-body fitting, flexible fitting, and de novo model building, as well as various combinations of these [3–6]. However, such approaches have primarily been applied to proteins, with less focus on bound small molecules. It may be particularly challenging to maintain the native intricacies of inter-molecular ligand-protein interactions during computational modeling. Recent efforts to combine physics-based docking with relatively low-resolution ligand densities [7–9] may be limited by reliance on an initial protein structure, which must be determined by separate methods, and substantially restricted from further refinement even in the case of overlapping protein and ligand densities. Flexible fitting methods can in principle refine a ligand and the surrounding protein pocket simultaneously, but typically require a masked ligand-only density, hindering automation. For most automated methods, successful modeling depends on a reasonably accurate initial model for the protein-ligand complex.

Recent artificial intelligence (AI)-based methods, including AlphaFold3 and its open-weights analogs such as Chai-1, offer new approaches to predict structures of protein-ligand complexes based on amino-acid sequences and ligand specification [10, 11]. Here we test the applicability of such a protocol, in combination with simulation-based flexible fitting where necessary, to support an automated strategy for modeling pharmaceutically relevant protein-ligand complexes into cryo-EM maps not represented in the AI training data (Fig. 1). For ten biomedically relevant targets, including cytosolic kinases, membrane-bound receptors and secondary transporters, ligand models generated in Chai-1 fit the target cryo-EM density with at least 82% accuracy relative to the deposited structure, either directly or after density-guided simulations. These results demonstrate the utility of combining generative AI with flexible fitting to automate ligand building in a variety of pertinent systems.

**Fig. 1.**
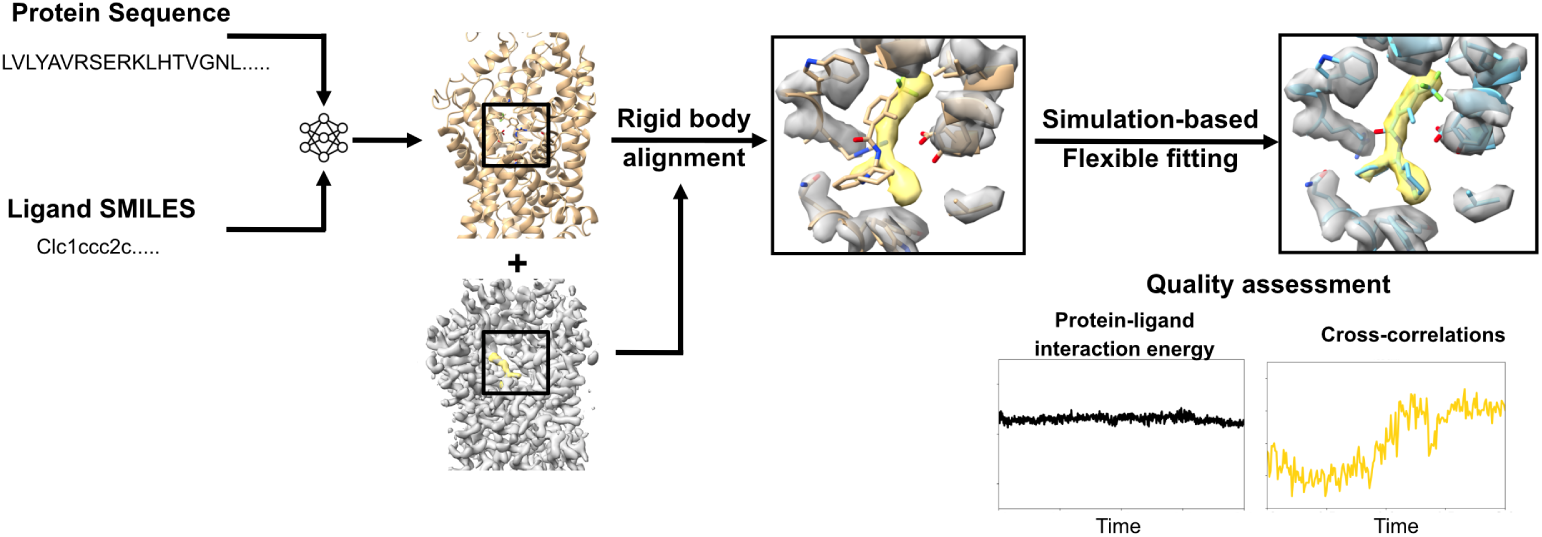
An approach to automated modeling of ligand-protein complexes. First, the protein sequence and ligand SMILES information were provided in Chai-1 to predict the protein-ligand complex structure. Then, rigid body alignment followed by molecular dynamics simulation-based flexible fitting were performed.

## Results

### An approach to automated modeling of ligand-protein complexes

To test the integration of generative AI with flexible fitting in building experimental structures of protein-ligand complexes, we first predicted five molecular models for each target using Chai-1 [11]. This open-weights model is based on comparable architecture and training strategies to those of AlphaFold3 [10], and has shown similar performance in predicting protein-ligand complexes [11]. For each target, we input the protein amino-acid sequence and a ligand specification using the simplified molecular input line-entry system (SMILES) (Fig. 1, left). The predicted complexes were then rigid-body aligned with their target cryo-EM maps using ChimeraX (Fig. 1, center) [12].

Next, we used density-guided molecular dynamics (MD) simulations in GROMACS [13] to fit the best Chai-1 model to the density (Fig. 1, right). Briefly, in this step, we applied additional forces to atoms of the protein and ligand scaled by the gradient of similarity between a simulated density based on the initial model and the reference cryo-EM map. No additional restraints were applied during these simulations, in order to enable conformational adjustments to both the protein and ligand during refinement. During fitting, we monitored the model-to-map cross-correlations (CCs) as a metric to track the quality of the fit, protein-ligand interaction energy (PLIE) for favorable interactions without clashes, and the generalized orientation-dependent all-atom potential (GOAP) score [14] for protein geometry. Notably, PLIE does not correspond to free energy which is a non-trivial parameter to estimate from our simulations. To minimize technical challenges arising from inconsistent ligand nomenclature, we focused on CCs to validate our fitted versus experimental structures (ground truth). Because a ground truth would not be available in fitting original data, information from the experimental structure was not used during model building, fitting or quality assessment, but served as a reference for validation.

### A test set of biomedically relevant protein-ligand complexes

We tested our approach on monomeric target complexes in the protein-ligand interactions dataset and evaluation resource (PLINDER) [15] (see Methods). Experimental structures for these test cases were reported after December 2021, such that they were not present in the Chai-1 training set. We filtered these for complexes containing native protein sequences under 1200 amino acids, along with pharmaceutically relevant small molecules (quantitative estimation of drug-likeness (QED) ≥ 0.7 [16]) within 4 Å of protein atoms. The resulting test set consisted of ten protein-ligand complexes of 400–1200 amino acids resolved to 2.7–3.7 Å, including two cytosolic kinases, two G protein-coupled receptors (GPCRs) and six secondary transporters (Table 1 and Fig. S1), and ligands with molecular weights from 218 to 442 g/mol (16-30 heavy atoms), 0-4 rotatable bonds, and 3-10 hydrogen bond donor/acceptor atoms (Table 2 and Fig. S2).

**Table 1.** Protein-ligand complexes tested in this study.

| Protein name | Ligand name | EMDB ID | Overall resolution (Å) | PDB ID | Extended sampling needed? |
| --- | --- | --- | --- | --- | --- |
| LRRK2 | MLi-2 | 41709 | 3.1 | 8TXZ [17] | X |
| PI3Kα | Alpelisib | 34271 | 2.7 | 8GUA [18] | X |
| H1R | Desloratadine | 38079 | 3.4 | 8X64 [19] | X |
| HCA3 | Acifran | 35447 | 3.2 | 8IHK [20] | X |
| GlyT1 | SSR504734 | 37494 | 3.2 | 8WFK [21] | X |
| CHT1 | Hemicholinium-3 | 36027 | 3.6 | 8J75 [22] | X |
| ThTr2 | Thiamine | 38691 | 3.7 | 8XV2 [23] | X |
| NET | Bupropion | 34722 | 3.0 | 8HFL [24] | X |
| OCT3 | Corticosterone | 14725 | 3.7 | 7ZH6 [25] | X |
| OATP1B1 | Estrone sulfate | 34911 | 3.2 | 8HND [26] | YES |

**Table 2.**
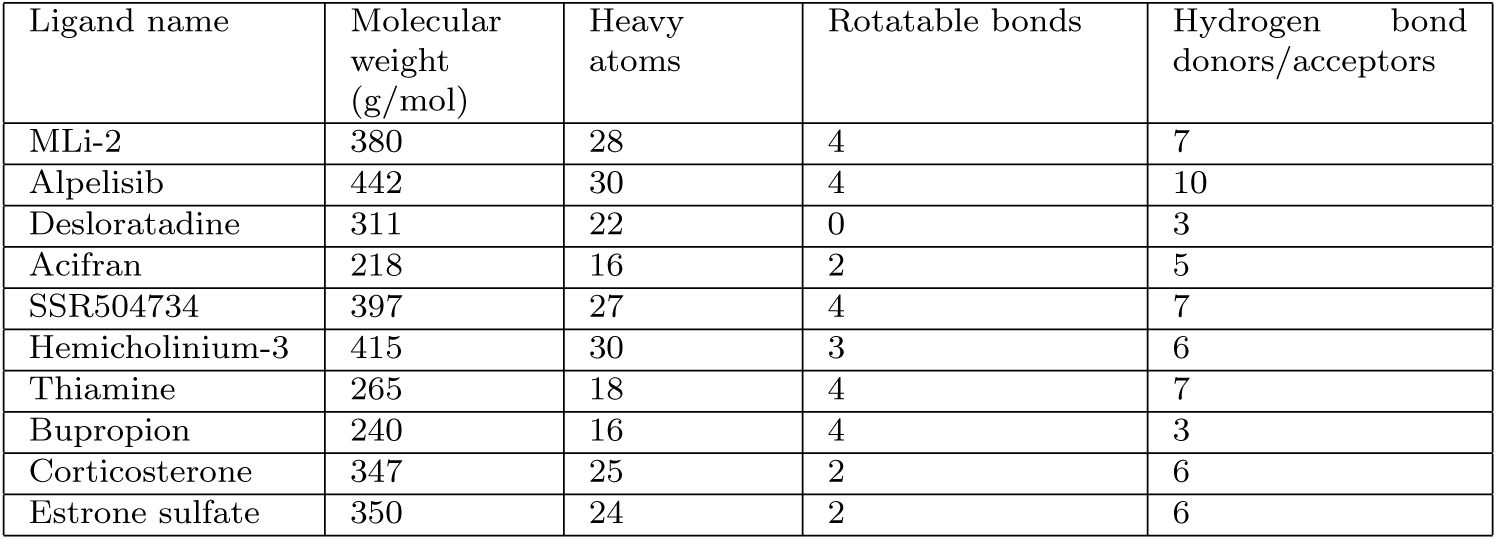
Properties of the ligands used in this study, extracted from PubChem. [37].

The kinases included 1) leucine-rich repeat kinase 2 (LRRK2), a multifunctional enzyme (1194 residues) and driver of heritable Parkinson’s disease [27]; and 2) a variant of phosphoinositide 3-kinase *α* (PI3K*α*), a lipid kinase of similar size (1096 residues) but distinct fold, mediating cellular growth signals in human cancers [28] (Table 1 and Fig. S1). The target complex with LRRK2 included MLi-2, an inhibitor with high affinity and selectivity [17]; the complex with PI3K*α* included BYL-719 (alpelisib), a Food and Drug Administration (FDA) approved compound for the treatment of solid tumors [29] (Table 2 and Fig. S2).

The membrane-bound receptors included 1) the histamine H1 receptor (H1R), a GPCR (438 residues) that is responsible for allergic and inflammatory symptoms [30]; and 2) the type-3 hydroxycarboxylic acid (HCA3) receptor, a smaller GPCR (387 residues) with only 13% identity to H1R, also involved in inflammation as well as neuroprotection [31] (Table 1 and Fig. S1). The target H1R complex included desloratadine, a second-generation antihistamine [19]; the complex with HCA3 included acifran, a drug capable of lowering plasma low-density lipoprotein concentrations [20] (Table 2 and Fig. S2).

The secondary transporters included 1) the sodium- and chloride-dependent glycine transporter 1 (GlyT1), a member of the solute carrier-6 (SLC6) family (652 residues) regulating both inhibitory and excitatory neurotransmission, and a target for the treatment of schizophrenia [21]; 2) the high-affinity choline transporter 1 (CHT1), part of the SLC5 family (580 residues) mediating reuptake of synaptic choline following neurotransmitter hydrolysis and involved in conditions such as depression and anxiety [32]; 3) the thiamine transporter 2 (ThTr2), a member of the SLC19 family (496 residues) that takes up dietary vitamins and is involved in diseases such as Wernicke’s encephalopathy [33]; 4) the norepinephrine transporter (NET), another SLC6 member (617 residues) which plays an essential role in the central nervous system and is a therapeutic target for emotional and cognitive disorders [34]; 5) the organic cation transporter 3 (OCT3), a member of the SLC22 family (556 residues) responsible for cellular uptake of cationic drugs in various tissues [35]; and 6) the organic anion transporting polypeptide 1B1 (OATP1B1), another SLC22 family member (691 residues) that transports a wide range of amphipathic organic anions, including clinical drugs [36] (Table 1 and Fig. S1). In these test cases, GlyT1 was bound to its inhibitor SR504734 [21]; CHT1 to its inhibitor hemicholinium-3 [22]; ThTr2 to its substrate thiamine [23]; NET to its inhibitor bupropion, an FDA-approved antidepressant [24]; OCT3 to its inhibitor corticosterone [25]; and OATP1B1 to its endogenous metabolite estrone sulfate [26] (Table 2 and Fig. S2).

### Imprecise prediction resolved by flexible fitting

For three of our test cases, including both kinases (LRRK2+MLi-2, PI3K*α*+alpelisib) and one GPCR (H1R+desloratadine), standard Chai-1 prediction followed by model-to-map rigid-body alignment generated ligand poses with at least 0.6 CC and -6 kJ/mol PLIE (Table 3). These poses corresponded to at least 90% ligand accuracy, as measured by CC relative to that of the ground-truth structure (see Methods) (Figs. 2 and S3). Indeed, although we report accuracy only for the best of five generated models, all predictions in these cases appeared largely consistent, deviating less than 0.1 in raw CC (Figs. S4 to S6). Protein structures were also similar to the ground truth, predicted with at least 88% accuracy for pocket residues within 4 Å of the ligand, and with at least 83% accuracy for the entire protein (Figs. S3 and S7). For these cases, although generative modeling appeared sufficient to approximate the target complexes as previously reported for other systems [11], we still performed flexible fitting and maintained a similar level of accuracy.

**Fig. 2.**
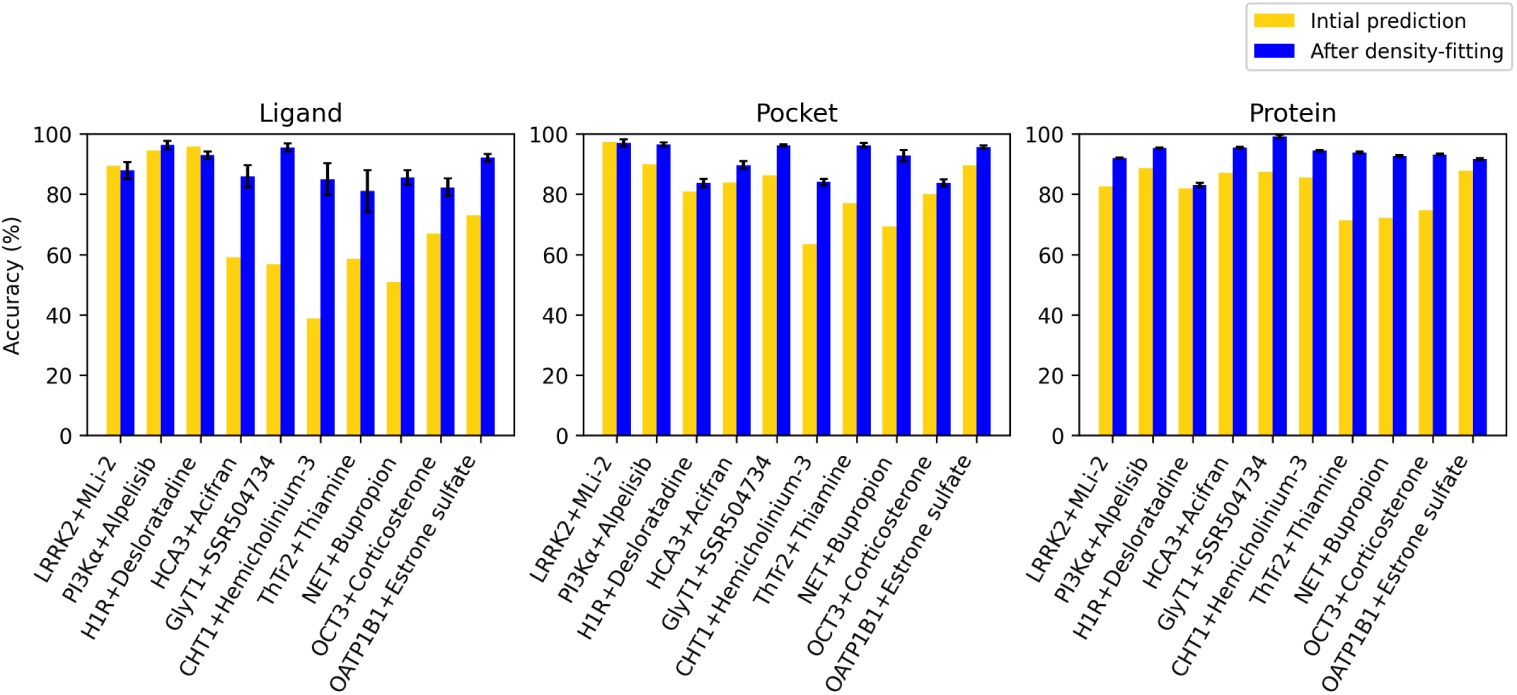
Accuracy of generative-AI models before and after flexible fitting in 10 test systems. Accuracy for ligand (left), pocket residues (center) and protein residues (right) based on model-to-map CC relative to that of the corresponding deposited structure, for the best initial generative-AI prediction (yellow) and after simulation-based fitting to the experimental map, when applied (blue). For fitted models, columns represent mean accuracy and standard error over the final 20 frames of the simulation.

**Table 3.** Protein-ligand interaction energies (PLIE) and model-to-map CCs for the best Chai-I prediction (Pred.), final complex from flexible fitting (Fitted) and ground truth based on the deposited experimental structure (GT) for each test system. PLIE values are scaled by number of ligand heavy atoms.

| System name | Ligand CC |  |  | Protein CC |  |  | Pocket CC |  |  | PLIE (kJ/mol) |  |
| --- | --- | --- | --- | --- | --- | --- | --- | --- | --- | --- | --- |
|  | Pred. | Fitted | GT | Pred. | Fitted | GT | Pred. | Fitted | GT | Pred. | Fitted |
| LRRK2+MLi-2 | 0.8 | 0.8 | 0.9 | 0.7 | 0.7 | 0.73 | 0.46 | 0.55 | 0.63 | -7.3 | -8.2 |
| PI3Kα+Alpelisib | 0.7 | 0.7 | 0.75 | 0.55 | 0.6 | 0.65 | 0.45 | 0.52 | 0.56 | -8.9 | -7.4 |
| H1R+Desloratadine | 0.64 | 0.62 | 0.68 | 0.52 | 0.55 | 0.71 | 0.52 | 0.54 | 0.7 | -11.7 | -7.3 |
| HCA3+Acifran | 0.35 | 0.61 | 0.76 | 0.52 | 0.6 | 0.71 | 0.54 | 0.53 | 0.67 | -8.9 | -6.6 |
| GlyT1+SSR504734 | 0.24 | 0.62 | 0.67 | 0.68 | 0.78 | 0.82 | 0.58 | 0.7 | 0.71 | -6.4 | -7.9 |
| CHT1+hemicholinium-3 | 0.2 | 0.7 | 0.8 | 0.43 | 0.64 | 0.8 | 0.53 | 0.61 | 0.67 | -12.6 | -10.1 |
| ThTr2+Thiamine | 0.4 | 0.7 | 0.8 | 0.5 | 0.7 | 0.72 | 0.51 | 0.74 | 0.8 | -15.8 | -20.1 |
| NET+Bupropion | 0.35 | 0.72 | 0.84 | 0.5 | 0.71 | 0.8 | 0.44 | 0.65 | 0.72 | -7.9 | -9.4 |
| OCT3+Corticosterone | 0.5 | 0.64 | 0.82 | 0.57 | 0.62 | 0.77 | 0.46 | 0.64 | 0.71 | -10.4 | -7.2 |
| OATP1B1+Estrone sulfate | 0.57 | 0.8 | 0.85 | 0.68 | 0.76 | 0.8 | 0.56 | 0.62 | 0.7 | -10.0 | -10.1 |

For the remaining seven test cases, ligand poses were predicted with less than 0.6 CC, corresponding to less than 71% accuracy and suggesting a need for further optimization. In three of these cases, including one GPCR (HCA3+acifran) and two transporters (GlyT1+SSR504734, CHT1+hemicholinium-3), standard Chai-1 generation followed by rigid-body alignment produced some of the least accurate ligand fits in this work (0.2-0.35 CC, or 40-60 % accuracy). For HCA3, the ligand only partly overlapped the target density in 1 of 5 generated models (Table 3 and Figs. 2, 3 and S8 to S10). However, overall protein conformation was predicted in these cases with at least 86 % accuracy (Table 3 and Figs. 2 and S11 to S13), including outward-facing states for both transporters (overall root mean squared deviation [RMSD], based on C*α* atoms, within 1.5 Å from their respective experimental structures [21, 22]), indicating that only the ligand fit needed substantial improvement. Indeed, density-guided MD simulations of each of these complexes for 2 ns improved ligand CC from 0.2-to 0.6-0.7 and accuracy from 40-60% to 85-95%. Accuracy also improved for the protein models, increasing from 63-86% to 84-96% for residues in the binding pockets, and 86-87% to 90-99% for the proteins overall. During our simulations, PLIE remained below -6 kJ/mol (Fig. 3). Similarly, GOAP scores largely remained below ground-truth values, indicating that geometric plausibility was retained (Fig. S14).

**Fig. 3.**
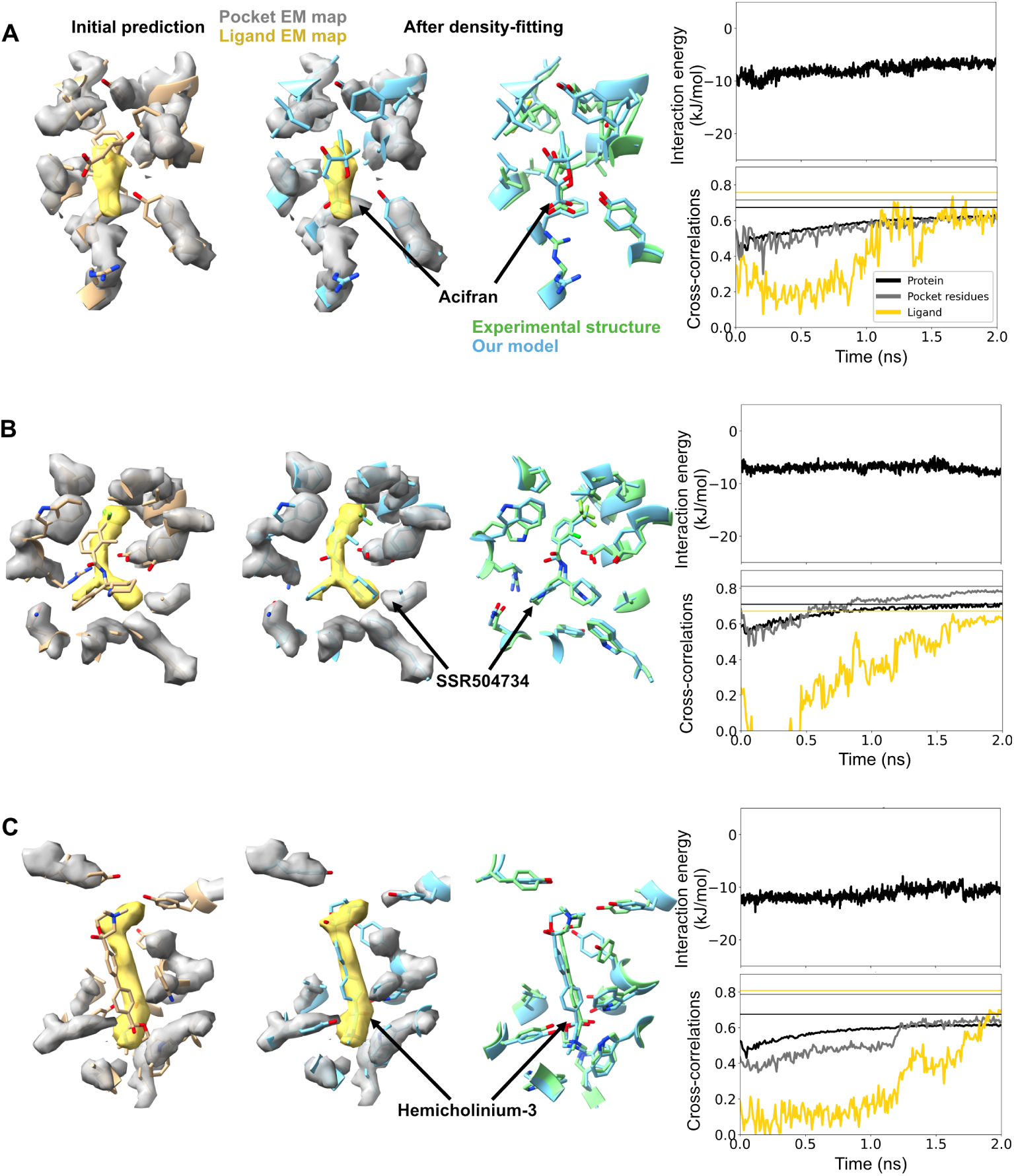
Imprecise prediction resolved by flexible fitting. (*Left*) Initial Chai-1 prediction of the binding pocket for the type-3 +hydroxycarboxylic acid receptor+acifran (A), glycine transporter+SSR504734 (B), and choline transporter+hemicholinium-3 (C) systems. Cryo-EM densities for the ligand (yellow) and pocket protein residues (silver) are shown in transparent. (*Middle*) Final frame of the density-fitting simulations (blue) along with the experimental structure (green). (*Right, top*) Time trace of the protein-ligand interaction energy is shown. (*Right, below*) Time trace of the cross-correlations of the ligand (yellow), pocket residues (silver), and the entire protein (black) during the density-fitting simulations. The vertical lines (with same color coding) represent values calculated using the experimental structure. Chai-1 prediction of all the 5 possible binding sites can be found in Figs. S8 to S10.

### Flexible fitting improves protein as well as ligand conformations

In three additional test cases, including the transporters ThTr2+thiamine, NET+bupropion, and OCT3+corticosterone, the protein conformations as well as ligand poses were predicted with relatively low accuracy (71-75% and 50-68% respectively) (Table 3 and Figs. 2, 4 and S15 to S17). Visual inspection indicated they deviated from the functional states assigned to the deposited structures, a particular challenge for targets like SLCs whose biological function involves conformational cycling [38]. Poor correspondence did not appear to correlate with the functional state or fold of the target density: ThTr2 and NET were resolved in inward-facing states [23, 24], while OCT3 was outward-facing [25], similar to GlyT1 and CHT1 in the previous section. ThTr2 and OCT3 are in the major facilitator superfamily, while NET as well as GlyT1 and CHT1 are in the LeuT family. Still, all best-fit models characterized in this work deviated less than 4 Å overall from the ground-truth structures deposited for their corresponding targets, such that density-guided simulations could be expected to accomplish reasonably accurate refinement [6]. Indeed, density-guided simulations improved accuracy of the overall protein models to 93-94%, and the ligands to 82-90%(Table 3 and Figs. 2 and 4), indicating that protein conformational changes as well as ligand fitting can be accommodated by this flexible fitting protocol. PLIE remained below -6 kJ/mol for all the three cases, indicating favorable packing between the ligand and protein.

**Fig. 4.**
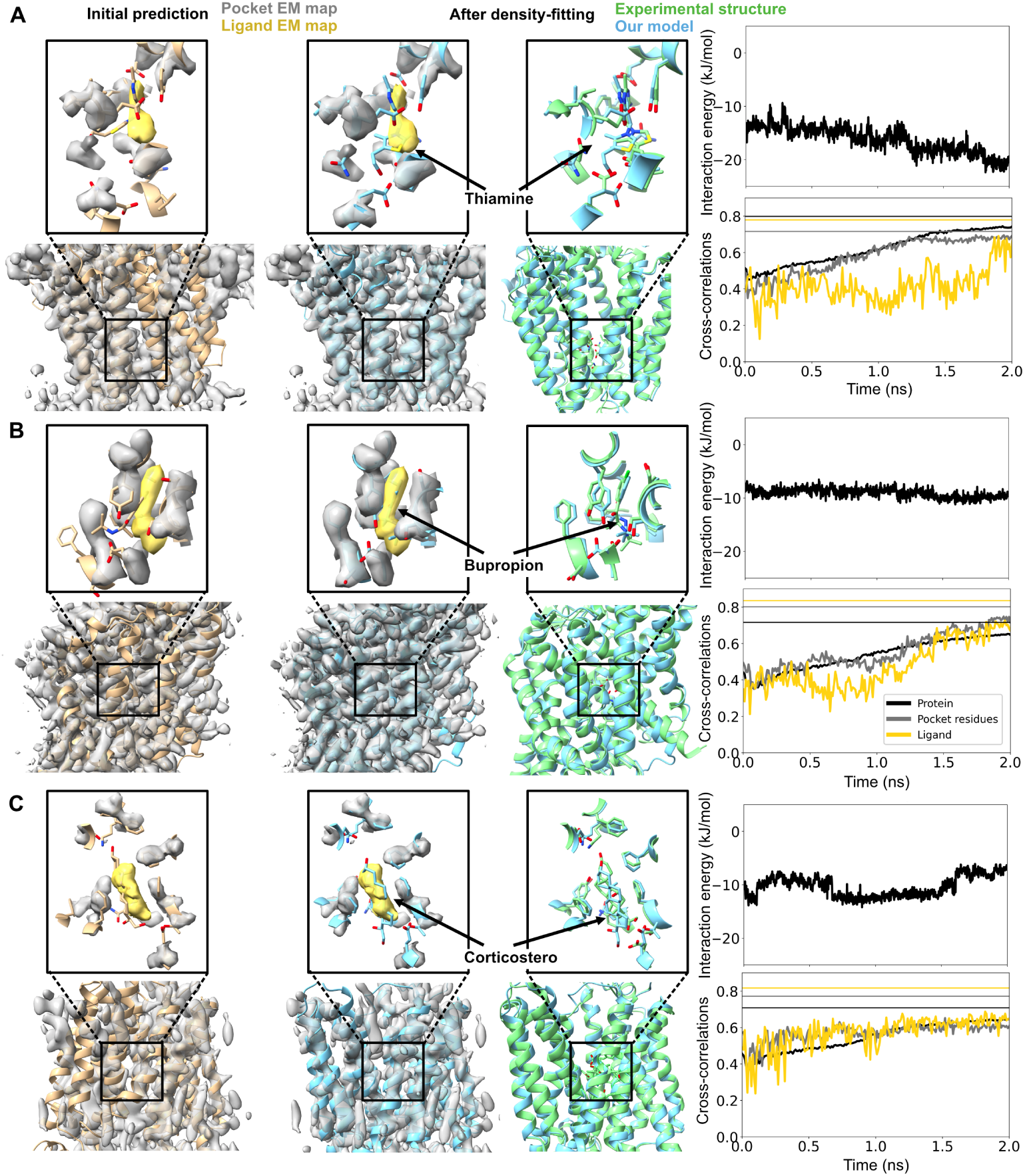
Flexible fitting improves protein as well as ligand conformations. (*Left*) Initial Chai-1 prediction of the binding pocket for the thiamine transporter 2+thiamine (A), norepinephrine transporter+bupropion (B), and organic cation transporter+corticosterone (C) systems. Same coloring scheme used as in Fig. 3. Alignment of the entire protein in addition to binding pockets are shown. (*Middle*) Final frame of the density-fitting simulations (blue) along with the experimental structure (green). (*Right, top*) Time trace of the protein-ligand interaction energy is shown.(*Right, bottom*) Time trace of the cross-correlations of the ligand (yellow), pocket residues (silver), and the entire protein (black) during the density-fitting simulations. The vertical lines (with same color coding) represent values calculated using the experimental structure. Chai-1 prediction of all the 5 possible binding sites can be found in Figs. S15 to S17.

### Challenging ligand poses improved by extended sampling in Chai-1

For the OATP1B1+estrone sulfate complex, standard Chai-1 prediction produced a best-fit model in the apparent outward-facing state (RMSD of 1 Å from the experimental target structure [26]) with 90% accuracy for the binding pocket, and 92% accuracy for the protein overall (Table 3 and Fig. 2,Fig. 5A,B). The ligand was modeled with 67% accuracy (Table 3 and Fig. 2), and improved to 77% after density-guided simulations. However, PLIE values above -1 kJ/mol during simulation suggested the ligand was not well coordinated in the binding pocket. Indeed, visual inspection indicated that even the best-fit ligand was flipped relative to the ground truth, a type of discrepancy that may not be easily corrected by flexible fitting (Fig. 5B). To improve the initial model, we generated an additional 20 predictions in Chai-1. The best resulting model included a ligand pose that was 71% accurate, and oriented similar to the experimental structure; ligand accuracy improved to 92% after density-guided simulations (Table 3 and Fig. 2,Fig. 5C,D). PLIE for this model was around -10 kJ/mol during the entire simulation. Thus, extended sampling in Chai-1 may usefully improve initial models for challenging ligand poses, although they were only required in one of our ten test cases.

**Fig. 5.**
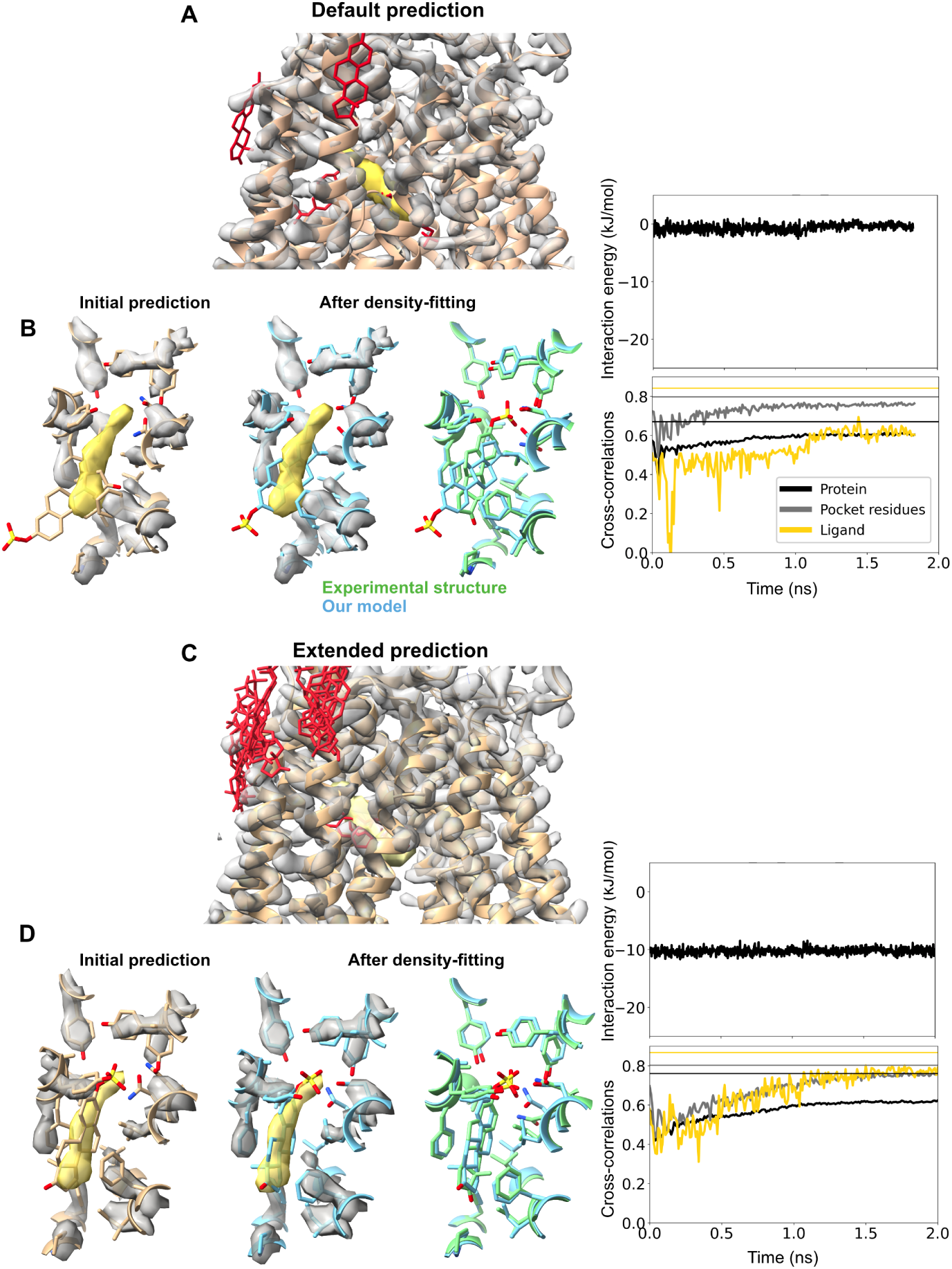
Challenging ligand poses improved by extended sampling in Chai-1. (A) Chai-1 prediction of 5 possible binding sites of estrone sulfate in a solute carrier organic anion transporting polypeptide, without pocket information. Same coloring scheme used as in Fig. 3. (B) One representative Chai-1 prediction of the binding pocket for complex, with the greatest cross-correlation for the ligand. Final frame of the density-fitting simulations (blue) along with the experimental structure (green). Time trace of the protein-ligand interaction energy, cross-correlations of the ligand, pocket residues, and the entire protein during the density-fitting simulations are shown. The simulation automatically terminated before reaching 2 ns, possibly due to the high force from the density. (C) Chai-1 prediction of 25 possible binding sites of the same complex. Same coloring scheme used as in Fig. 3. (D) One representative Chai-1 prediction of the binding pocket for complex, with the greatest cross-correlation for the ligand. Simulations time traces are shown as in the panel B.

## Discussion

Here, we showcase how AI models can complement MD simulations to automatically and accurately fit small molecules into cryo-EM maps, limiting user effort and bias during modeling. When benchmarked against ten biomedically relevant entries from the EMDB, our approach fits ligands with 82-95% accuracy relative to deposited structures. The resulting models would consitute appropriate templates for final manual or automated refinement, or in some cases for direct analysis and deposition.

Our approach addresses several challenges in automated structure determination of protein-ligand complexes [9, 39–41]. First, the flexible fitting protocol implemented here enables refinement of the protein alongside the ligand; the protein structure does not need to be accurately built in advance to achieve an accurate model of the complex. Second, the generative AI step requires only the amino acid sequence and SMILES specification as inputs, avoiding any need for structural templates. The initial models thus generated also do not require rebuilding of unresolved atoms, residues, loops, or other regions, as is often necessary when preparing experimental structures for MD simulations. Third, our method does not require knowledge of the ligand-binding location or coordinating residues, a nontrivial inference when studying novel complexes. It is plausible that this approach may also be applicable to simultaneously model multiple ligands, including ions and cofactors, as allowed by both AlphaFold3-like models and density-guided simulations [10, 11].

As noted in Fig. 4, generative AI methods might not predict the relevant conformational state of a protein-ligand complex. This may be a particular concern for proteins such as transporters that undergo conformational changes between functional states. The SLCs investigated in this study operate by various alternating-access mechanisms, cycling between inward- and outward-facing states. Proteins in the LeuT superfamily such as GlyT1, CHT1, and NET generally employ gated-pore mechanisms, while members of the major facilitator superfamily such as ThTr2, OCT3, and OATP1B1 are associated with rocker-switch mechanisms respectively [38]. Possibly due to the diversity of relevant structure for each of these transporters, AI methods can fail to predict a given state. Nonetheless, conformational differences in all cases studied here could be accommodated with reasonable accuracy by flexible fitting. We previously reported that transporters for which functional states deviate more than 4 Å benefit from the generation and clustering of an ensemble of initial models [42], an approach that could also prove valuable in more challenging cases of ligand fitting. As AI predictions continue to improve, the success of flexible fitting is likely to extend to a broader range of systems.

A potential limitation of our approach is its performance with proteins that lack evolutionary data or have highly flexible structures. Previous studies have shown that AlphaFold2 can struggle to predict new protein conformations in such scenarios [43]. This challenge underscores the importance of the density-guided molecular dynamics step in our approach, as it enables a good fit even when none of the generated models closely matches the target conformation. Incorporating previously described multi-step approaches, such as iterative density-guided simulations with progressively increasing resolution or enhanced sampling techniques [39], could further improve our pipeline performance in more challenging cases. Another limitation of our approach is that forcefields for small molecules in classical MD simulations may not be highly accurate. However, given the relatively high force arising from the cryo-EM density potential, dependence on classical forcefields should be minimized. Future developments in AI models may further circumvent this problem by improving initial ligand positioning.

## Methods

### Data curation

To curate test cases, we used the recently released protein-ligand interaction structural database called PLINDER [15]. Using PLINDER’s parquet file format, we extracted recently released (January, 2022- June, 2024) cryo-EM monomeric protein ligand complexes that contains drug-like small molecules (with quantitative estimation of drug-likeness [QED] value ≥ 0.7 [16]). The date cutoff ensures that our test cases were not in the training set of Chai-1 (cutoff date: Dec, 1st, 2021) [11]. This resulted into a dataset of 23 test cases, which was narrowed down to 10 after removing proteins with unnatural chimeric constructions, nonconclusive ligand binding sites (judged based on no atoms being closer than 4 Å in the binding pocket), homologous pairs, proteins with more than 1200 amino acids.

### Chai-1 predictions

We used the Chai-1 webserver lab.chaidiscovery.com to predict our protein-ligand complexes, where the protein amino acid sequence and ligand SMILE string were used as input [11]. The webserver by default outputs 5 predictions. For the case of solute carrier organic anion transporting polypeptide bound to estrone sulfate, 20 additional predictions were generated.

### System preparations

Each protein-ligand complexes predicted by Chai-1 was prepared for molecular dynamics (MD) simulations using the CHARMM-GUI [44] webserver. Each system was solvated with TIP3P water [45] and neutralizing in 0.15 M KCl to generate systems containing in the range from 90,000 to 210,000 atoms. It was shown previously that the membrane does not substantially improve density guided simulation results [46], so we did not include membrane building step into the pipeline for simplicity. The systems were energy minimized and then relaxed in simulations at constant pressure (1 bar) and temperature (310K) for 1 ns, during which the position restraints on the protein and ligands were gradually released. The restraints were used as recommended by CHARMM-GUI.

### Density-guided simulations

Density-guided MD simulations in this study were performed using GROMACS-2024 [13] utilizing CHARMM36m [47] and CHARMM General Force Field (CGenFF) [48] force field parameters for proteins and ligands, respectively. Bonded and short-range nonbonded interactions were calculated every 2 fs, and periodic boundary conditions were employed in all three dimensions. The particle mesh Ewald (PME) method [49] was used to calculate long-range electrostatic interactions with a grid spacing below 0.1 nm*^−^*^3^. A force-based smoothing function was employed for pairwise non-bonded interactions at 1 nm with a cutoff of 1.2 nm. Pairs of atoms whose interactions were evaluated were searched and updated every 20 steps. A cut- off of 1.2 nm was applied to search for the interacting atom pairs. Constant pressure was maintained at 1 bar using the c-rescale barostat[50] and temperature was kept at 310K with the v-rescale thermostat [51]. Forces from density-guided simulations were applied every N = 2 steps. The scaling factor for density-guided simulation forces of 10^3^ kJ/mol was combined with adaptive force scaling.

### Analysis

Cross-correlations to the target map during simulations were calculated using the “mdff check -ccc” command in Visual Molecular Dynamics (VMD) [52]. Protein-ligand interaction energies were calculated using the “gmx energy” feature in GROMACS [13]. System visualizations were carried out by ChimeraX [12]. Structure quality check during MD simulations were done using the generalised orientation-dependent all-atom potential (GOAP) score matrix, where low values indicate better structure [14]. Model accuracy is calculate using the following formulae:

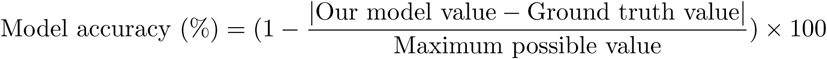

## Acknowledgments

We thank Dr. Marta Bonaccorsi, Dr. Stavros Azinas and the Molecular Biophysics Stockholm environment for valuable feedback and discussion. Computational resources were provided by the National Academic Infrastructure for Supercomputing in Sweden (NAISS 2024/3-49).

## Data and code availability

All data needed to evaluate the conclusions are present in the paper or the Supplementary Information. The generated models, density-guided simulation input files and trajectories can be found on Zenodo (doi 10.5281/zenodo.14842872). The simulation method is available as a component in the GROMACS molecular simulation toolkit.

## Author Contributions

Conceptualization: NH Data curation: NH

Formal analysis: NH Funding acquisition: NH, EL Investigation: NH Methodology: NH

Project administration: RJH, EL Resources: NH, EL

Software: NH Supervision: RJH, EL Validation: NH Visualization: NH

Writing – original draft: NH

Writing – review & editing: NH, RJH, EL

