## Supplementary materials for "Cryo-EM ligand building using generative AI and molecular dynamics"

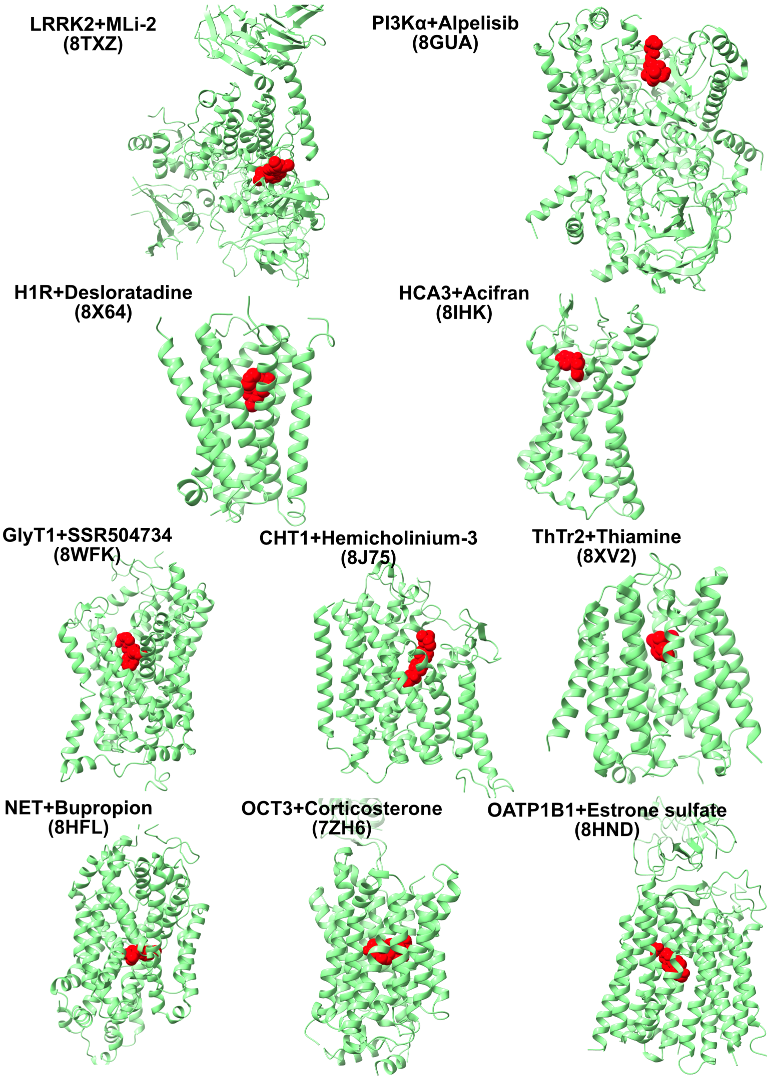


**Fig. S1** Experimental structures of the proteins tested in this study. PDB IDs are described in brackets after the names of each protein. Red sphere shows the ligands bound to the proteins.

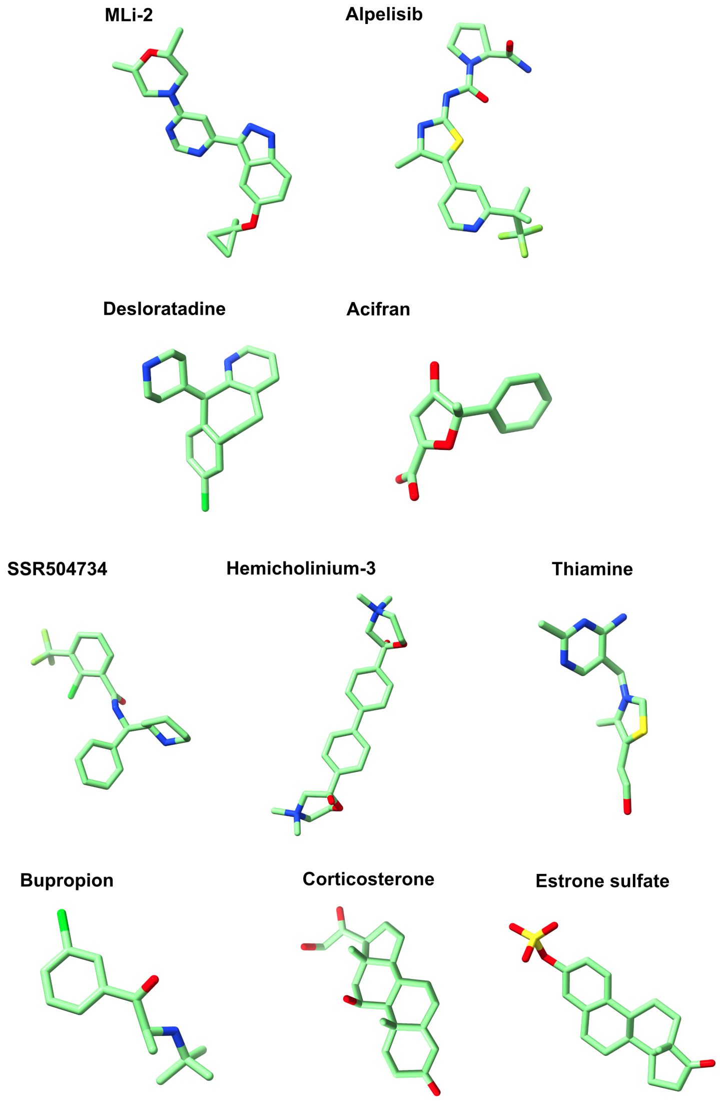


**Fig. S2** Ligand overview used in this study.

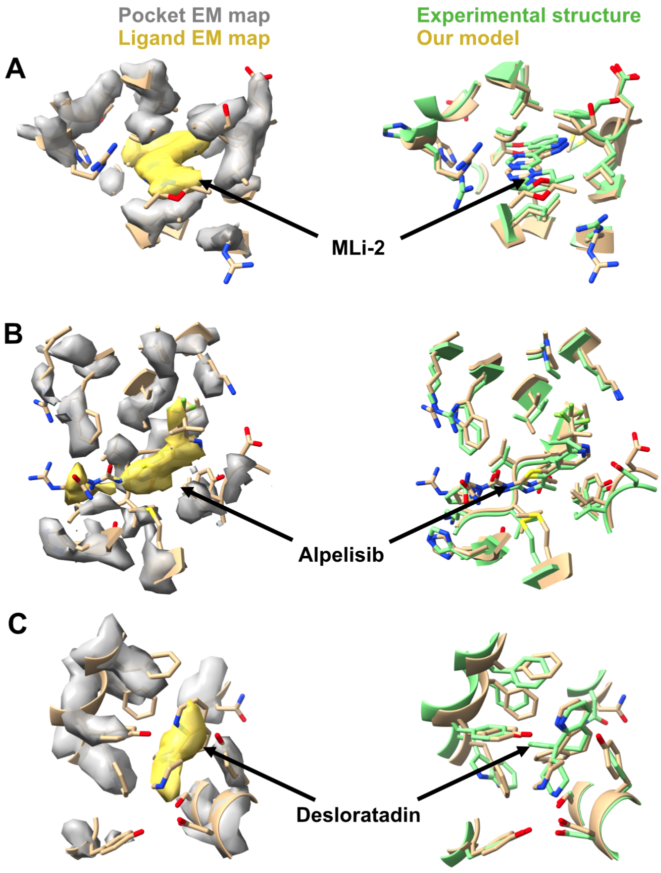


**Fig. S3 Accurate prediction of protein-ligand complexes.** (*Left*) Initial Chai-1 prediction of the binding pocket for the, leucine-rich repeat kinase 2+MLi-2 (A), phosphoinositide 3-kinase $\alpha$+alpelisib (B), and histamine H1 receptor+desloratadine (C) systems. Cryo-EM densities for the ligand (yellow) and pocket protein residues (silver) are shown in transparent. (*Middle*) Predicted structure (yellow) along with the experimental structure (green). Alignments were done using the entire protein in ChimeraX Matchmaker module. Chai-1 prediction of all the 5 possible binding sites can be found in Figs. S4 to S6.

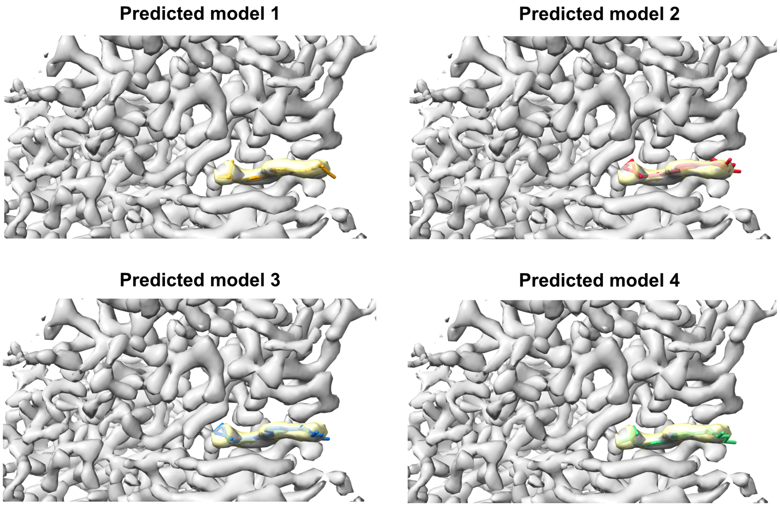


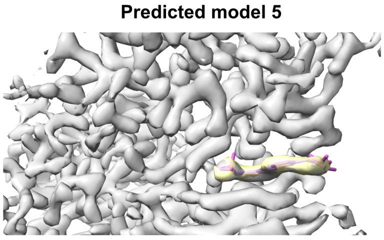


| Predicted model | Cross-correlations |
|---|---|
| 1 | 0.80 |
| 2 | 0.78 |
| 3 | 0.85 |
| 4 | 0.80 |
| 5 | 0.82 |

**Fig. S4** Chai-1 prediction of 5 possible binding sites of MLi-2 in a leucine-rich repeat kinase 2. Cryo-EM densities for the ligand (yellow) and pocket protein residues (silver) are shown in transparent. Cross-correlation values for the ligands are shown in the table below.

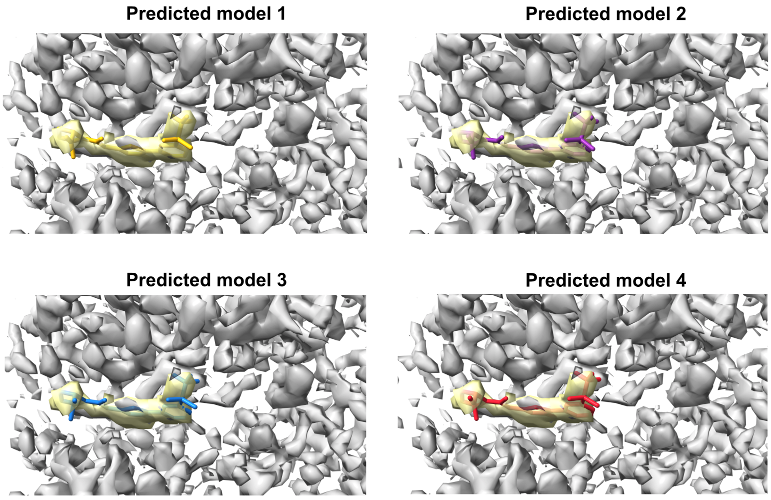


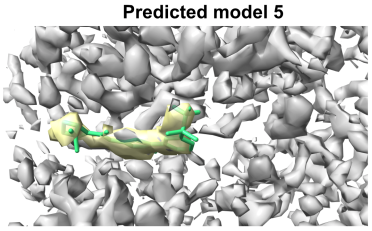


| Predicted model | Cross-correlations |
|---|---|
| 1 | 0.72 |
| 2 | 0.69 |
| 3 | 0.67 |
| 4 | 0.66 |
| 5 | 0.63 |

**Fig. S5** Chai-1 prediction of 5 possible binding sites of alpelisib in a phosphoinositide 3-kinase $\alpha$. Cryo-EM densities for the ligand (yellow) and pocket protein residues (silver) are shown in transparent. Cross-correlation values for the ligands are shown in the table below.

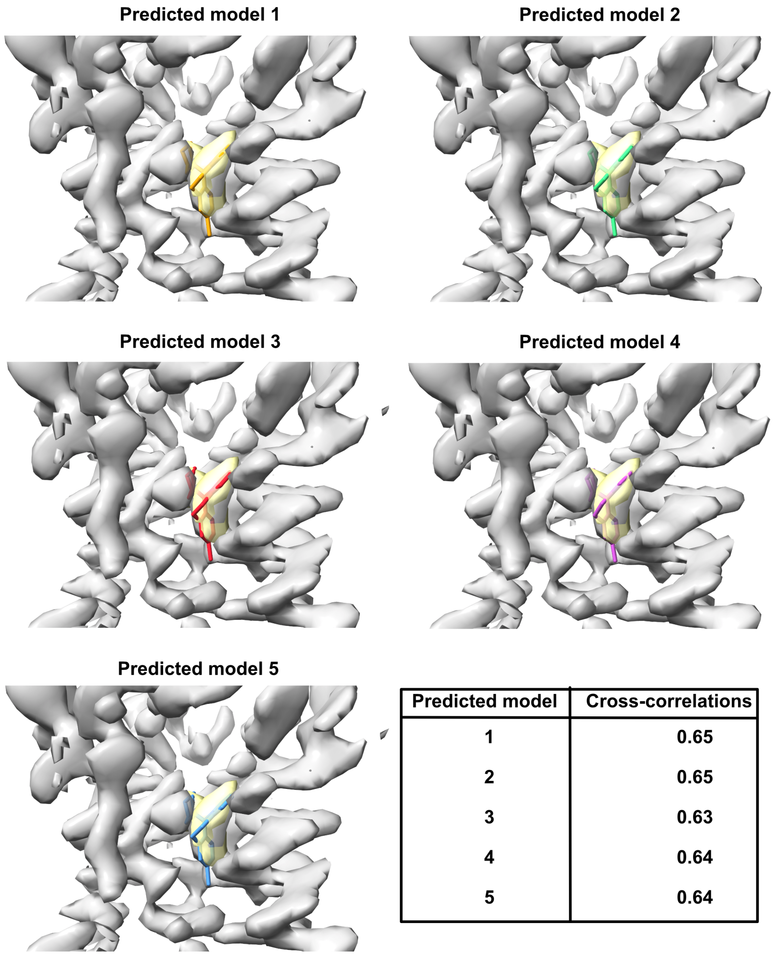


| Predicted model | Cross-correlations |
|---|---|
| 1 | 0.65 |
| 2 | 0.65 |
| 3 | 0.63 |
| 4 | 0.64 |
| 5 | 0.64 |

**Fig. S6** Chai-1 prediction of 5 possible binding sites of desloratadine in a histamine H1 receptor. Cryo-EM densities for the ligand (yellow) and pocket protein residues (silver) are shown in transparent. Cross-correlation values for the ligands are shown in the table below.

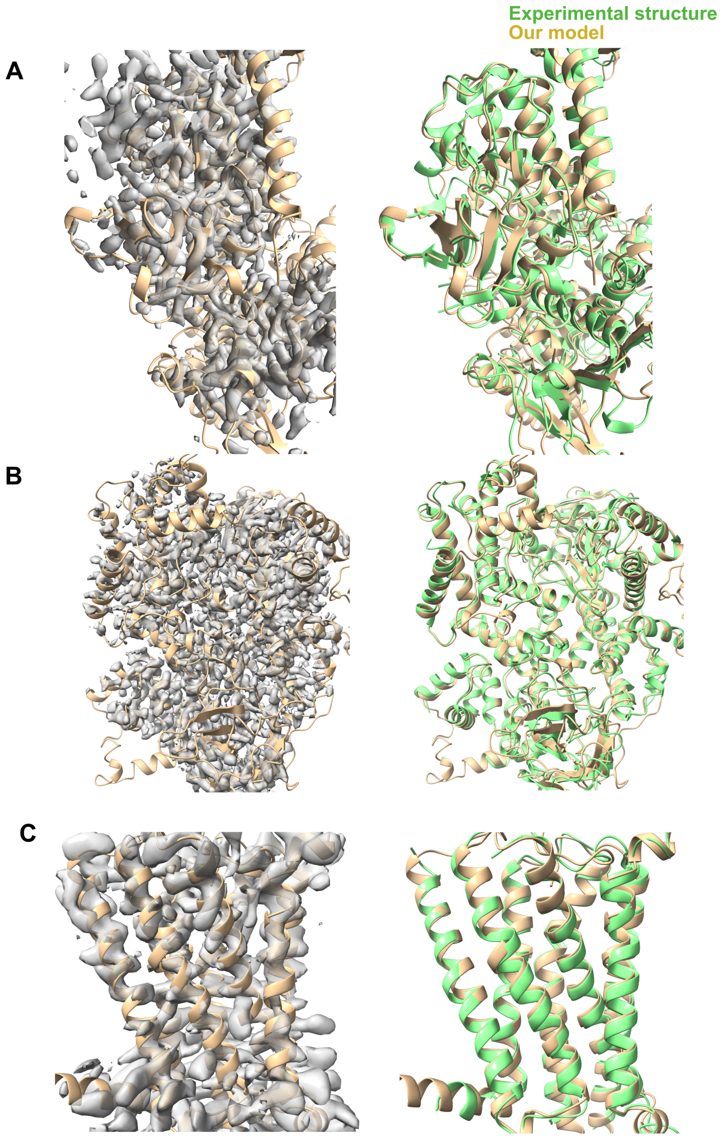


**Fig. S7** Chai-1 prediction of 3 complexes, shown in (Fig. S3), focusing on the entire protein:leucine-rich repeat kinase 2+MLi-2 (A), phosphoinositide 3-kinase $\alpha$+alpelisib (B), and histamine H1 receptor+desloratadine (C) systems. Predicted structure (yellow) along with the experimental structure (green) are shown on the right.

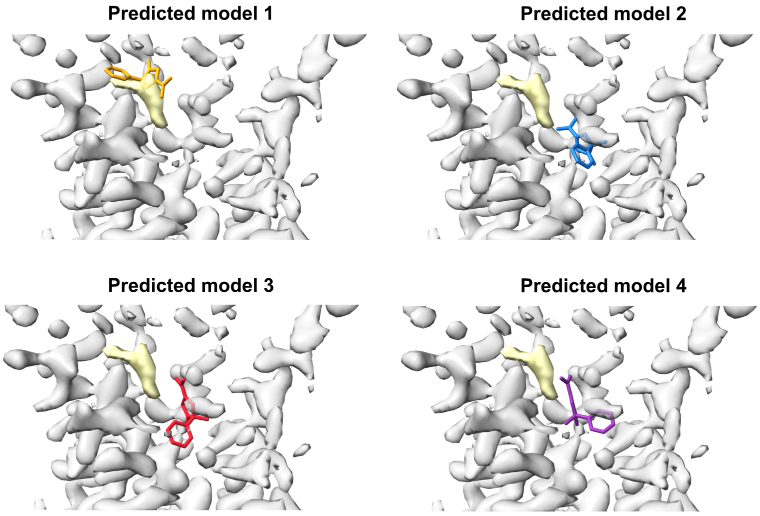


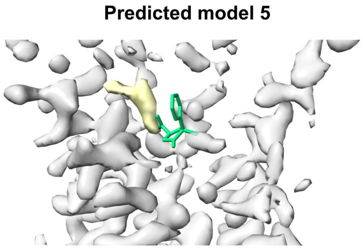


| Predicted model | Cross-correlations |
|---|---|
| 1 | 0.36 |
| 2 | 0.0 |
| 3 | 0.0 |
| 4 | 0.0 |
| 5 | 0.0 |

**Fig. S8** Chai-1 prediction of 5 possible binding sites of acifran in a type-3 hydroxycarboxylic acid receptor. Cryo-EM densities for the ligand (yellow) and pocket protein residues (silver) are shown in transparent. Cross-correlation values for the ligands are shown in the table below.

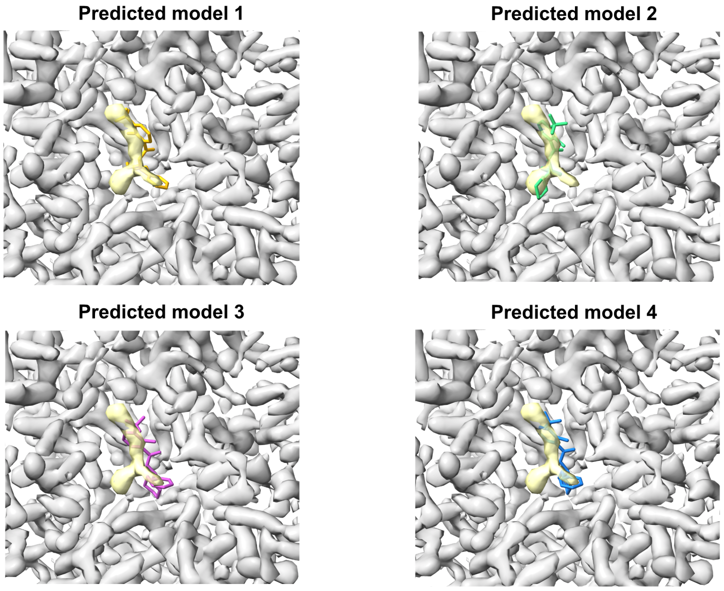


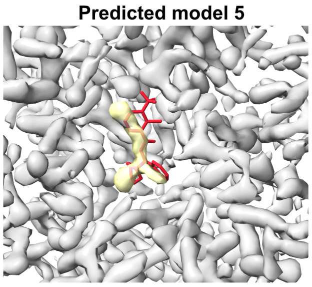


| Predicted model | Cross-correlations |
|---|---|
| 1 | 0.24 |
| 2 | 0.28 |
| 3 | 0.08 |
| 4 | 0.14 |
| 5 | 0.01 |

**Fig. S9** Chai-1 prediction of 5 possible binding sites of SSR504734 in a glycine transporter. Cryo-EM densities for the ligand (yellow) and pocket protein residues (silver) are shown in transparent. Cross-correlation values for the ligands are shown in the table below.

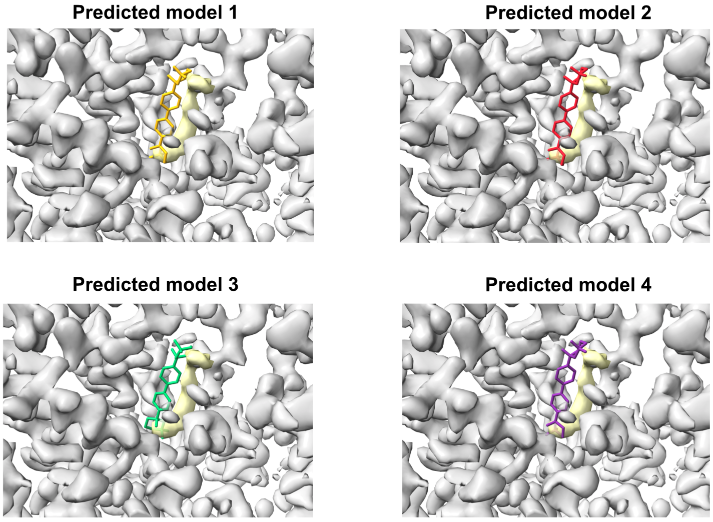


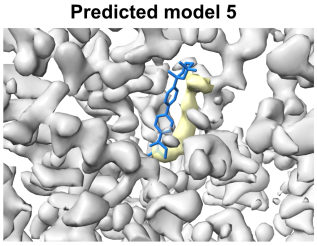


| Predicted model | Cross-correlations |
|---|---|
| 1 | 0.2 |
| 2 | 0.22 |
| 3 | 0.22 |
| 4 | 0.18 |
| 5 | 0.17 |

**Fig. S10** Chai-1 prediction of 5 possible binding sites of hemicholinium-3 in a choline transporter. Cryo-EM densities for the ligand (yellow) and pocket protein residues (silver) are shown in transparent. Cross-correlation values for the ligands are shown in the table below.

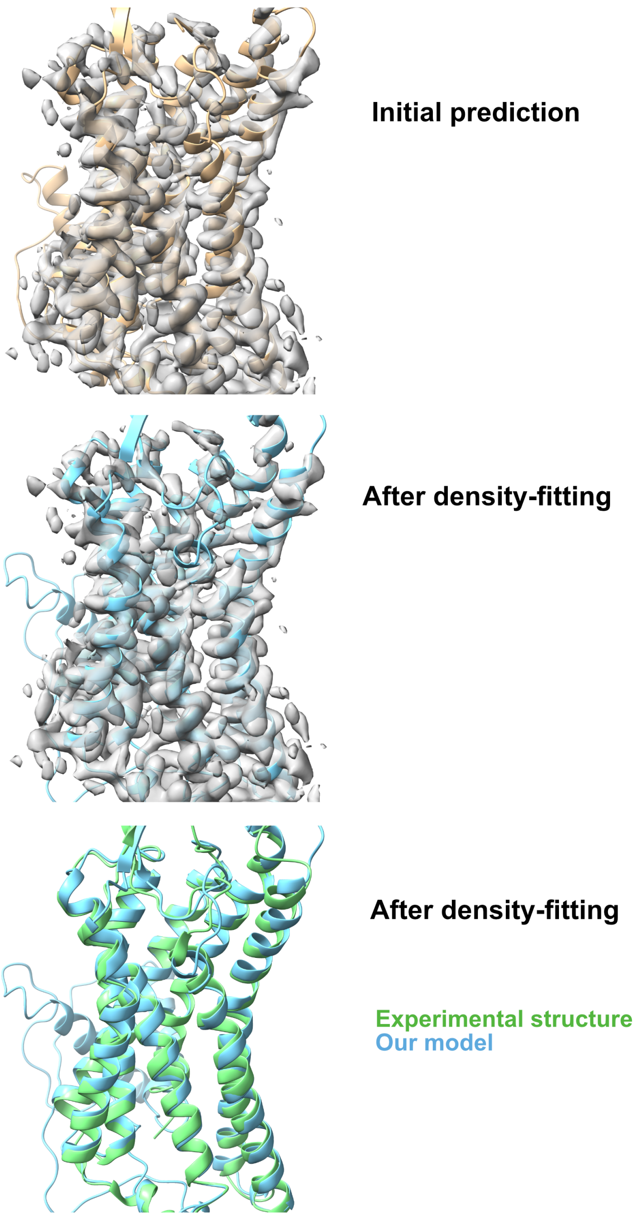


**Fig. S11** Initial Chai-1 prediction (yellow), final frame of the density-fitting simulations (blue) along with the experimental structure (green) of the type-3 hydroxycarboxylic acid receptor+acifran complex.

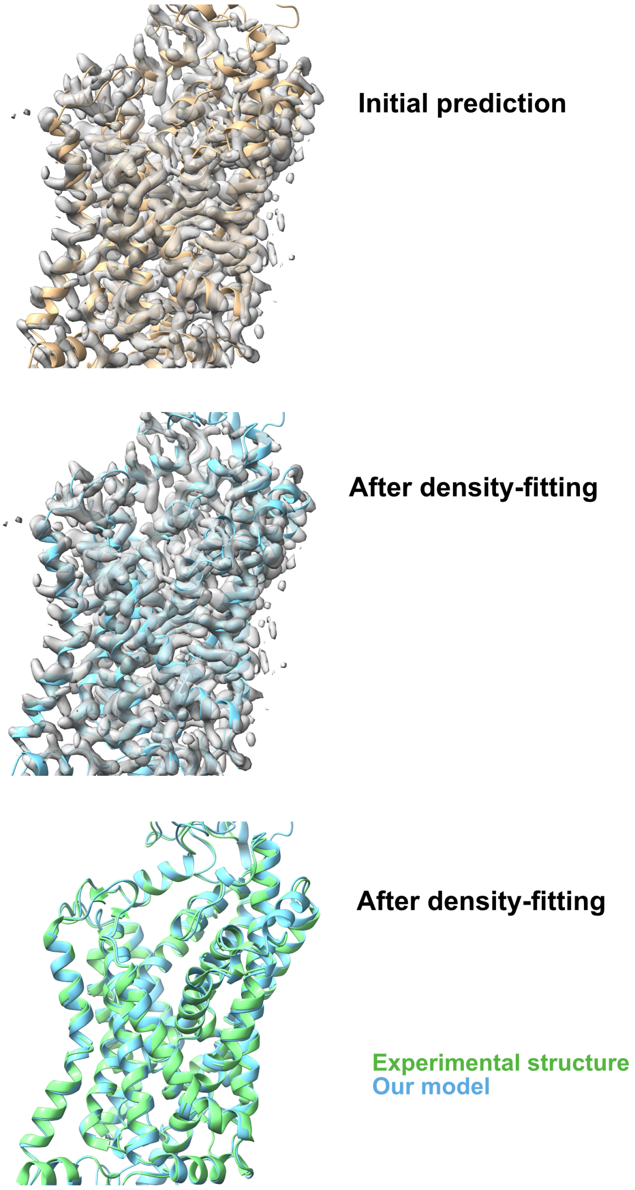


**Fig. S12** Initial Chai-1 prediction (yellow), final frame of the density-fitting simulations (blue) along with the experimental structure (green) of the glycine transporter 1+SSR504734 complex.

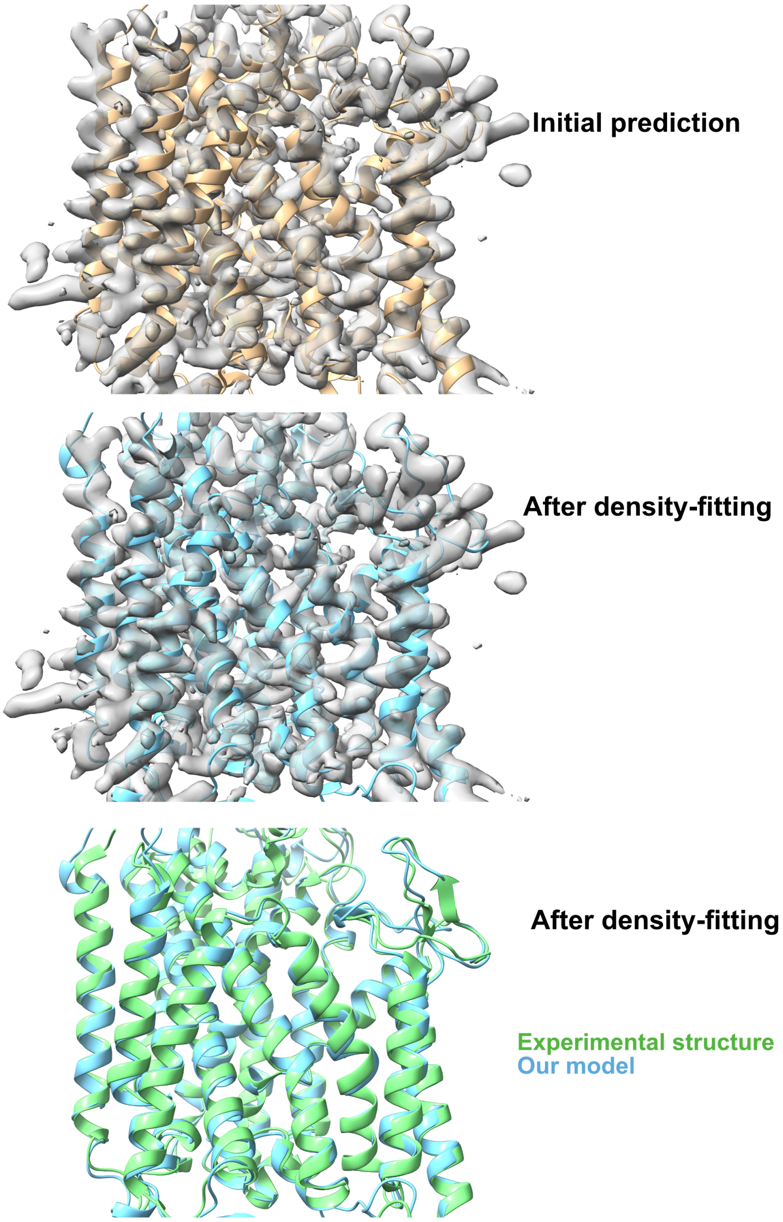


**Fig. S13** Initial Chai-1 prediction (yellow), final frame of the density-fitting simulations (blue) along with the experimental structure (green) of the choline transporter+hemicholinium-3 complex.

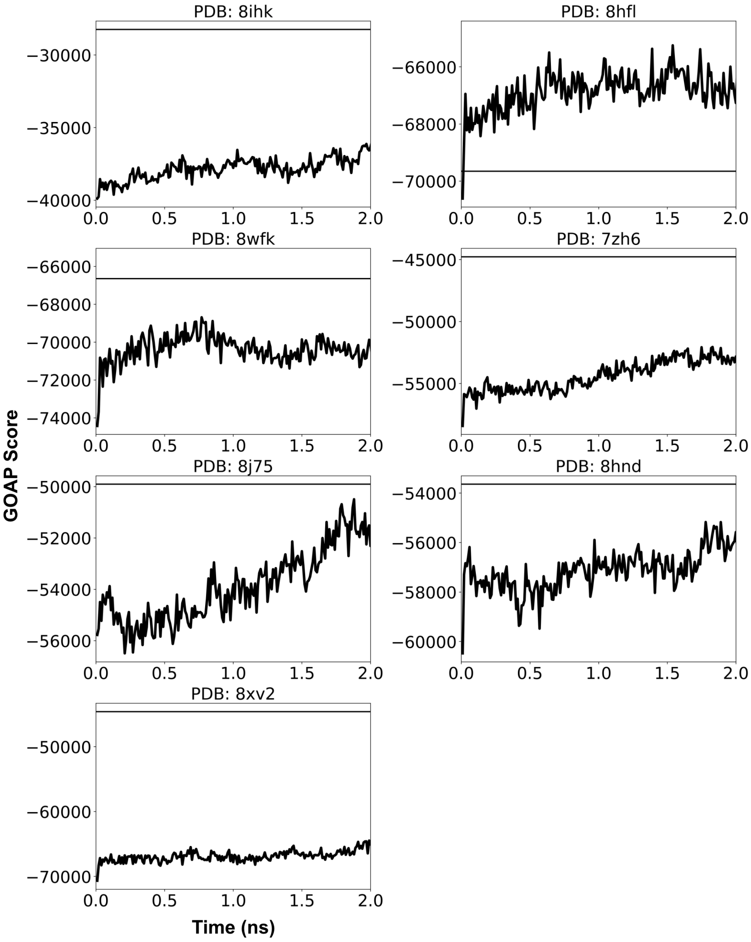


**Fig. S14** GOAP score calculation of the entire protein during our simulation. GOAP score for the experimental structure is depicted in the horizontal line. Lower the GOAP score, better the structure quality of the protein.

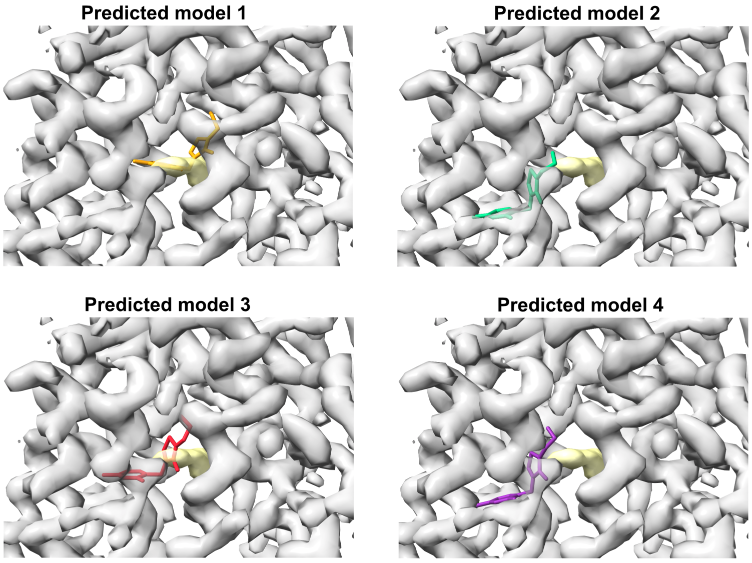


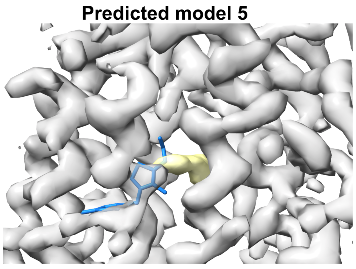


| Predicted model | Cross-correlations |
|---|---|
| 1 | 0.37 |
| 2 | 0.04 |
| 3 | 0.19 |
| 4 | 0.04 |
| 5 | 0.05 |

**Fig. S15** Chai-1 prediction of 5 possible binding sites of thiamine in a thiamine transporter 2. Cryo-EM densities for the ligand (yellow) and pocket protein residues (silver) are shown in transparent. Cross-correlation values for the ligands are shown in the table below.

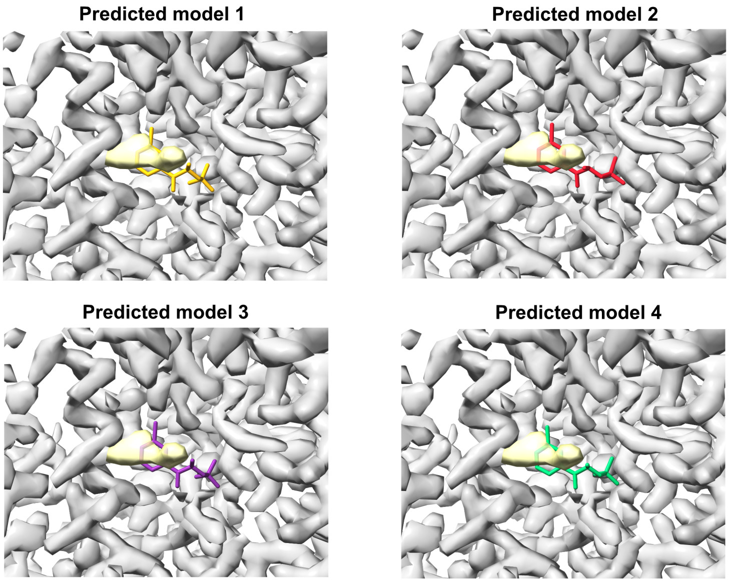


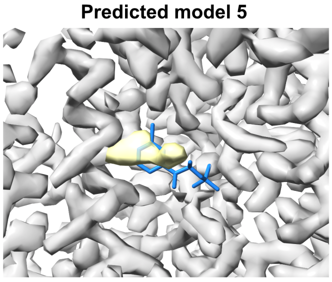


| Predicted model | Cross-correlations |
|---|---|
| 1 | 0.34 |
| 2 | 0.25 |
| 3 | 0.29 |
| 4 | 0.30 |
| 5 | 0.30 |

**Fig. S16** Chai-1 prediction of 5 possible binding sites of bupropion in a norepinephrine transporter. Cryo-EM densities for the ligand (yellow) and pocket protein residues (silver) are shown in transparent. Cross-correlation values for the ligands are shown in the table below.

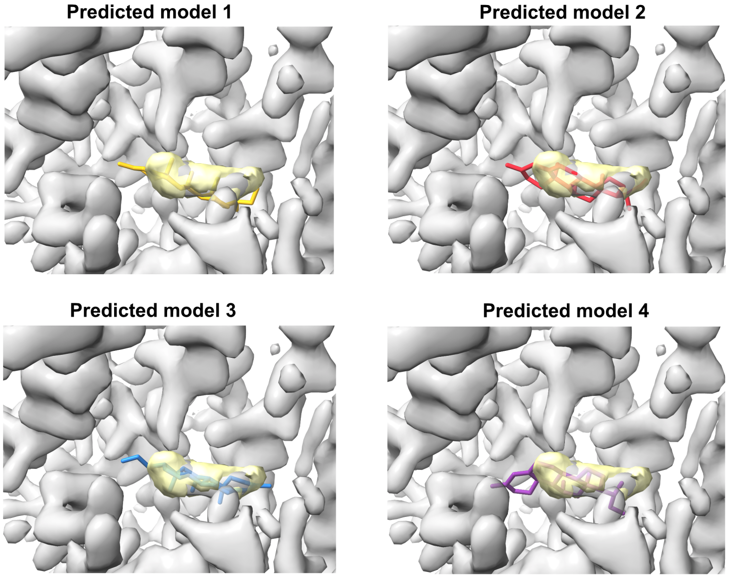


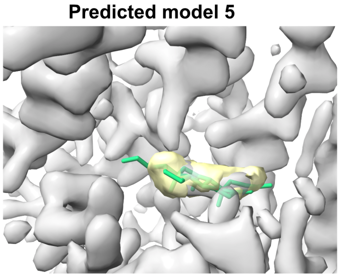


| Predicted model | Cross-correlations |
|---|---|
| 1 | 0.50 |
| 2 | 0.70 |
| 3 | 0.68 |
| 4 | 0.60 |
| 5 | 0.70 |

**Fig. S17** Chai-1 prediction of 5 possible binding sites of corticosterone in an organic cation transporter. Cryo-EM densities for the ligand (yellow) and pocket protein residues (silver) are shown in transparent. Cross-correlation values for the ligands are shown in the table below.
